# TRACE: Open-Source Software for Quantifying Somatic Variation of Tandem Repeats by Capillary Electrophoresis

**DOI:** 10.64898/2026.01.22.701167

**Authors:** Andrew Jiang, Kevin Correia, Tammy Gillis, Esaria L. Oliver, Benjamin P. Jones, Branduff McAllister, Alan Mejia Maza, Marcy E. MacDonald, Ricardo Mouro Pinto, Vanessa C. Wheeler, James F. Gusella, Zachariah L. McLean

**Affiliations:** Molecular Neurogenetics Unit, Center for Genomic Medicine, Massachusetts General Hospital, Boston, MA, 02114, USA; Department of Neurology, Harvard Medical School, Boston, MA, 02115, USA; Medical and Population Genetics Program, the Broad Institute of M.I.T. and Harvard, Cambridge, MA, 02142, USA; Department of Genetics, Blavatnik Institute, Harvard Medical School, Boston, MA, 02115, USA

## Abstract

Expanded short tandem DNA repeats are implicated in over 60 human disorders. In many, somatic instability (SI) of the repeat plays a critical role in disease pathogenesis. For example, SI in vulnerable neurons is a key driver of clinical symptoms in Huntington’s disease. Quantifying SI has traditionally relied on PCR followed by capillary electrophoresis, with metrics describing the shape of repeat size distributions, such as the expansion index. However, current tools often require costly proprietary software, are time-consuming, and rely on custom pipelines that vary between labs. To address these challenges, we developed Tandem Repeats Analysis by Capillary Electrophoresis (TRACE), an open-source software that processes fragment analysis data end-to-end, from raw files to SI metrics. Additionally, we created an associated web app TRACE-shiny (https://traceshiny.mgh.harvard.edu/), for interactive usage. Outputs from TRACE benchmarked against published datasets confirm its utility for studying genetic and pharmacological modifiers of SI. TRACE eliminates the need for proprietary software or custom pipelines, making advanced tools for analysis of somatic repeat expansion widely accessible.

## INTRODUCTION

Expanded short tandem repeats (STRs) in DNA are implicated in over 60 human disorders, a number that continues to grow with advancements in sequencing technologies (Chen et al., 2025; Depienne & Mandel, 2021). In many STR disorders, somatic instability (SI) of the repeat plays a critical role in disease pathogenesis. For example, in Huntington’s disease (HD), CAG repeat expansions in vulnerable neurons are somatically unstable (Handsaker et al., 2025; Mätlik et al., 2024; Swami et al., 2009), and genetic association studies implicate DNA repair genes in modifying this expansion (Genetic Modifiers of Huntington’s Disease, 2025), directly linking somatic repeat instability to clinical symptoms. With growing academic and therapeutic interest activities in this area across >60 STRs, tools for quantifying SI metrics are increasingly necessary to enable automation, reproducibility, and high-throughput data analysis.

Quantifying SI has traditionally relied on polymerization chain reaction amplification (PCR) followed by capillary electrophoresis, using metrics that describe the shape of repeat distributions. The instability index was devised to quantify the mean change in CAG length relative to the inherited repeat length (Lee et al., 2010). This can be broken down into the contraction or expansion components, with the later become widely adopted for quantifying the average repeat increase in mice or human tissues (Mouro Pinto et al., 2020; Mouro Pinto et al., 2025). However, existing tools for processing raw data often depend on expensive proprietary software that require manual processing, such as GeneMapper (Applied Biosystems). Alternatively, researchers have assembled custom pipelines (McAllister et al., 2022) that incorporate tools like the Fragman R package (Covarrubias-Pazaran et al., 2016). While effective, these tools were not designed specifically for STR analysis, often require extensive tuning, and do not correct run-to-run variation. Such *ad hoc* solutions lead to variability between labs and are challenging to adapt or customize for specific SI applications.

To address these challenges and enhance throughput and reproducibility, we present the Tandem Repeats by Capillary Electrophoresis (TRACE) R package. This includes new algorithms for raw capillary electrophoresis data processing and batch correction, along with unique features for SI experiments. We also developed TRACE-shiny (https://traceshiny.mgh.harvard.edu/), a web-based platform for code-free interfacing with the TRACE package. TRACE and TRACE-shiny eliminates the need for proprietary software or custom pipelines, making advanced tools for consistent analysis of somatic repeat expansion widely accessible.

## MATERIAL AND METHODS

### TRACE R package

The underlying code for processing fragment analysis data and generation of metrics is built upon our R package TRACE (https://zachariahmclean.github.io/trace/). This aggregates our algorithms described previously (Genetic Modifiers of Huntington’s Disease, 2025; McLean et al., 2024) into a unified framework. Users interact primarily with the high-level function trace(), which processes raw fragment analysis data and produces structured output. This output can then be supplied to calculate_instability_metrics() to generate metrics and visualizations of the data. The pipeline is built on custom R classes and is designed to work efficiently with the TRACE-Shiny application.

### Processing signal, fitting ladder and assigning base pair

The FSA files (Applied Biosystems fragment analysis file format with.fsa extension) are imported into memory in R using read.abif from the seqinr package (Charif & Lobry, 2007). To fit the ladder, we test combinations of observed peaks for their linear correlation to the ladder. However, testing all possible combinations becomes computationally prohibitive as the number of identified peaks increases relative to the ladder. We developed an optimization to reduce the total number of combinations based on the assumption that simultaneous testing of all possible combinations is unnecessary. Since ladder peaks follow a predictable size-ordered pattern, we can process reference sizes sequentially in chunks (Supplementary Figure 1). When assigning smaller reference sizes, we only consider combinations up to a point where the remaining observed peaks would all line up with the remaining number of ladders. This eliminates vast numbers of irrelevant combinations from consideration, while also accounting for the non-linearity of capillary electrophoresis data (Elder & Southern, 1983). For each chunk of reference sizes, we test combinations from a constrained window of candidate observed peaks (the remaining observed peaks would fit one-to-one with the remaining size standard), progressively moving this window as we assign larger reference sizes. At each chunk, the algorithm branches by a user-defined top-N, with non-promising branches pruned by an R-squared threshold. Base-pair sizes are interpolated using a generalized additive model with cubic regression splines using the mgcv package (Wood, 2010), which provides a flexible, accurate sizing model by accounting for slight non-linearities in the data. Peaks are identified in the ladder and trace data with findpeaks from pracma (https://CRAN.R-project.org/package=pracma).

### Allele calling

In this software, metrics are calculated from a single allele. This simplifies analysis in many samples, such as i) mouse models with expanded repeats in the context of human gene sequence, where there is a single amplifiable repeat allele, such *Hdh*^Q111^ (Q111) (Wheeler et al., 1999) and R6/2 (Menalled & Chesselet, 2002) models, or ii) human cell models where the expanded repeat of interest is typically >100 units larger than the normal allele(s), such that the normal allele(s) can be ignored (Ferguson et al., 2024; McLean et al., 2024). For HD human samples with alleles in the inherited adult-onset range, multiple alleles can be identified. But for simplicity, metrics are only calculated from the longer of the two alleles. The modal allele is identified within each sample by clustering peaks into “peak regions” based on height and size thresholds, then calling the tallest in the cluster as the main allele.

### Repeat calling

Repeat sizes for each peak are estimated based on the size of the PCR product, the number of nucleotides in the repeat unit, and the number of non-repeat nucleotides flanking the repeat. We intentionally do not report repeat sizes as integers, since they are derived from continuous raw signal with base-pair positions predicted relative to a size ladder. Additionally, exact integer calls are unnecessary for calculating instability metrics, so repeat sizes are therefore retained as continuous values to avoid inaccuracies introduced by rounding.

Since repeat-containing amplicons migrate non-linearly relative to the ladder, their base-pair sizes may be underestimated and show batch-to-batch variation. To address this issue, we also provide several important features including algorithms for correcting batch effects through a simple correction or by accurate repeat sizing. The simple batch correction algorithm lines up common samples across runs and adds a correction factor to address run-to-run variability. First, a smoothing algorithm is applied to locate the most representative modal size. Second, batch effects are estimated via mixed modeling using the lme4 package (Bates et al., 2015), providing flexibility for unbalanced data and handling non-overlapping sample sets. For accurate repeat sizing, a predefined repeat length provided in the metadata is assigned to the modal allele and adjacent repeat units. This creates a linear model for the relationship between base-pair size and repeat length. For both batch and repeat correction, a correction factor is applied to the peaks of all samples.

To evaluate the accuracy of the repeat-calling algorithm, we compared the modal allele called in samples from ABI 3130XL Genetic Analyzer processed manually using GeneMapper (v5.0) with those processed by TRACE. A total of 872 HD knock-in Q111 mouse tissue samples across 32 fragment analysis runs were analyzed (Supplementary Figure 2). Each fragment analysis submission had a sample with a known, validated repeat length for best practice accurate repeat sizing. For the GeneMapper analysis, bins of repeat sizes of *HTT* CAG-containing PCR amplicons, that incremented by one CAG repeat, were manually adjusted to line up with the repeat length of the modal peak of the validated reference sample. For processing the data in TRACE, the run and sample information was recorded in the metadata file using the categories “batch_run_id” (date of fragment analysis run), “batch_sample_id” (id of the validated reference sample that allows linking across runs for quality control), and “batch_sample_modal_repeat” (repeat length of the modal peak of the validated reference sample) and the “correction” parameter was set to “repeat”.

The underestimation of repeat size and the spacing between consecutive peaks was calculated using the same dataset as above. To calculate the spacing in base pairs between consecutive peaks (Supplementary Figure 2A), for each sample, the data were subsetted to peaks ranging from 110 to 130 to select only robust peaks and the median taken of the difference between consecutive peaks. This metric helps demonstrate the consistent narrow difference between peaks which should theoretically be exactly three base pairs. Next, the modal peak underestimation for each sample was found by dividing the repeat length calculated directly from base-pair size to the repeat length accurately determined with TRACE. This helps quantify the overall shortfall in modal repeat length when relying just on base-pair size. For Supplementary Figure 2B, the median value for each metric across samples was computed to enable correlation across runs. This helps demonstrate how the batch-to-batch differences in spacing between consecutive peaks might help explain the variability of allele repeat length.

### Instability metrics and benchmarking

The main instability metrics of interest are calculated as described previously (Lee et al., 2010) but also explained briefly below. To help with clarity, we introduced a new term “index peak”, defined as the reference peak within a fragment analysis trace. This may correspond to the inherited repeat length in a mouse or the time zero sample in a cell culture experiment. It is primarily used for instability index and related calculations and may or may not be the modal peak in a trace.

The expansion ratio (also known as peak proportional sum or somatic expansion ratio) focuses on the proportion of signal contributed by expansion or contraction peaks relative to the index peak, making it less sensitive to noise and well-suited for analyzing human samples with adult-onset ranges (Genetic Modifiers of Huntington’s Disease, 2025; Lee et al., 2019). We benchmarked the expansion ratio from the TRACE software to a previous pipeline we recently reported for 733 HD blood samples of the Registry dataset (Genetic Modifiers of Huntington’s Disease, 2025) with patients with adult onset repeat lengths.

The expansion index as originally described (Lee et al., 2010) measures expanded repeat lengths relative to the index peak, weighted by peak signal and distance from the index peak. It provides insight into the spread or diversity of repeat sizes by finding the average repeat change and is predominantly used for samples where there are mixed cell populations with differing rates of repeat expansion, which can produce a bimodal distribution of non-repeat expanding cells and expanding cells (e.g., mouse disease model brain tissue). For benchmarking, we reanalyzed FSA files from 879 Q111 mouse (with a single expanded CAG repeat) tissues across 32 fragment analysis runs, with a 5% relative peak height threshold and the TRACE parameter “force_whole_repeat_units” and accurate repeat sizing as described above, to match the repeat level GeneMapper data.

Average repeat change calculates the difference in weighted mean repeat size between the sample and the index samples (McLean et al., 2024). This metric is particularly intuitive for studying population-level shifts in cell models, where repeat distributions often remain approximately normally distributed. It is almost identical to the “instability index change” (instability index of the sample subtracted by the instability index of the index sample) (Ferguson et al., 2024), which uses the instability index formula rather than weighted mean, so relies more on accurate index peak assignment. We compared the average repeat change from the TRACE software to values generated with a custom pipeline that included FSA file processing with GeneMapper (McLean et al., 2024). These 53 samples were from the RPE1-AAVS1-CAG115 cell model testing the effect of cell division and dox-induced transcription.

Finally, the contraction index is the inverse of the expansion index (Lee et al., 2010) and measures the contracted repeat lengths relative to the index peak, weighted by peak signal and distance from the index peak. For benchmarking, we used 138 samples with a CCCTCT repeat associated with X-linked dystonia parkinsonism (XDP), with samples taken from various regions in post-mortem brain tissue (Mejia Maza et al., 2026).

### TRACE web server

We developed a standalone web-server application, TRACE-shiny (https://traceshiny.mgh.harvard.edu). TRACE-shiny was built using the shiny package in R (https://CRAN.R-project.org/package=shiny) and leverages shiny reactivity and plotly (Shalabh, 2021) visualizations to interactively guide the user through the fragment analysis pipeline. The focal point of the web server is to enable users without experience in R to progress through the TRACE pipeline. The interactivity element of TRACE-shiny enables additional features including 1) The ability to dynamically adjust ladder peaks that were incorrectly assigned (see *processing raw signal*). 2) Interactive assignment of the index repeat used in instability metrics calculations. 3) Tailored plotting tools designed to enhance visualization and emphasize major differences between raw signal traces. These include the option to normalize traces between samples by the highest peak or the sum of all peaks. The raw data can also be aggregated and the average found using best fit curve by local polynomial regression (loess).

## RESULTS

Here we present R package TRACE and associated web server TRACE-shiny. The TRACE R package provides functions in R for processing and analyzing raw fragment analysis data (FSA files) (Figure 1). The TRACE-shiny web server adds a user-friendly interface to guide analysis without previous R programming knowledge. TRACE-shiny enhances the analysis with a layer of interactivity and provides quickly accessible summary data at each step of the analysis. Below we highlight some of the unique features within this software and provide benchmarking of repeat instability metrics across a range of contexts.

**Figure 1.**
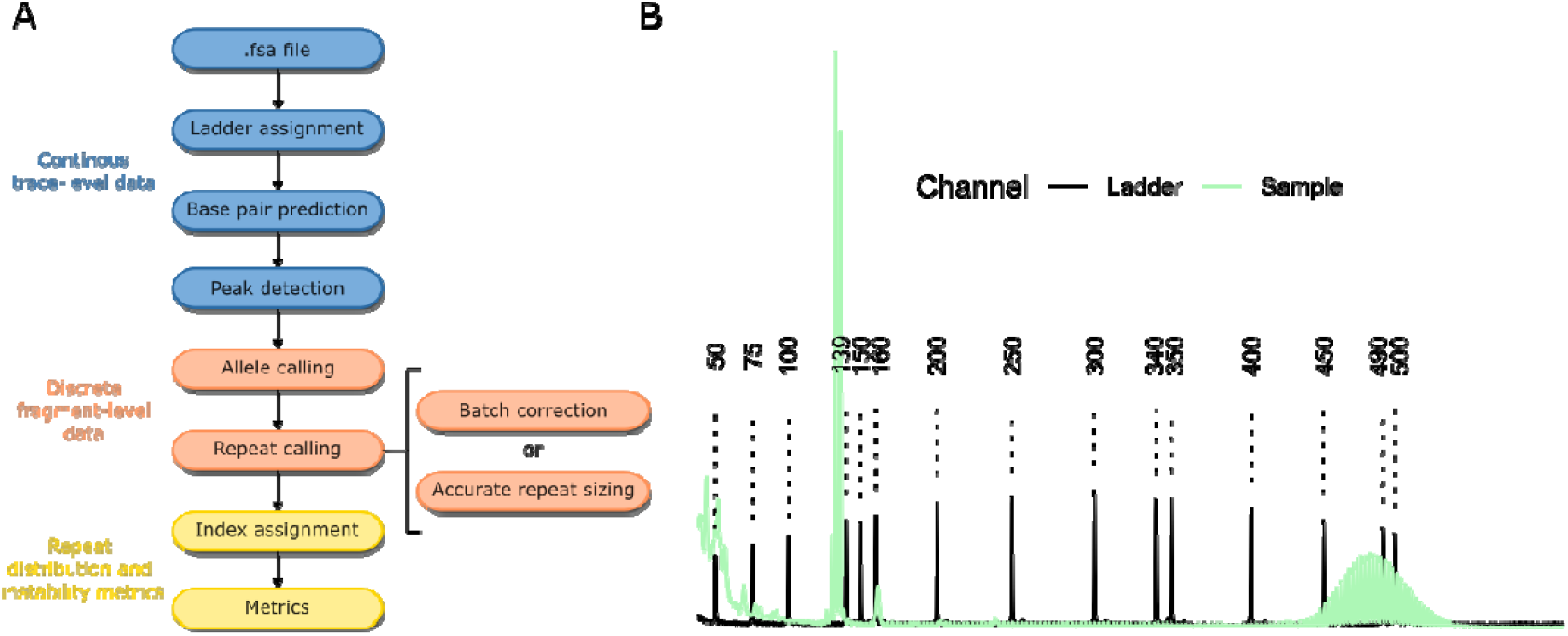
Data processing pipeline. (A) pipeline underlying the TRACE R package to process FSA files through to instability metrics. (B) An example of the raw signal for the size ladder (black) and sample (green).

### Ladder assignment algorithm

A significant challenge in capillary electrophoresis is assigning the ladder of known base-pair sizes to the corresponding peaks in the raw signal. In TRACE, we test combinations of observed peaks for their fit to the ladder using an optimization that minimizes the total number of combinations to test. Briefly, the observed ladder peaks are tested for linear fit in consecutive chunks, which simultaneously overcomes non-linearity of capillary electrophoresis data (Southern, 1979) while constraining combinations to prevent fruitless searches where insufficient peaks remain to be assigned. We tested this approach using challenging ladders that have contamination into the ladder channel, which introduces more peaks than are usually expected and makes the assignment difficult by combinatorics (Supplementary figure 3). The TRACE approach reduces the total number of combinations to test by 10-1000-fold, depending on the sample and window size used (figure 2A). The accuracy also depends on the window size used, with window sizes too small or large resulting in incorrect assignments (figure 2A). In our tests using this challenging dataset, this approach reduced the time from a mean of 21 seconds per sample to test all possible combinations to 0.07 seconds with a window size of five. To prevent the algorithm from getting trapped in local optimal fits, we implemented branching at each step, which only moderately increased combinations but was required for correct ladder fitting in several samples (figure 2B).

**Figure 2.**
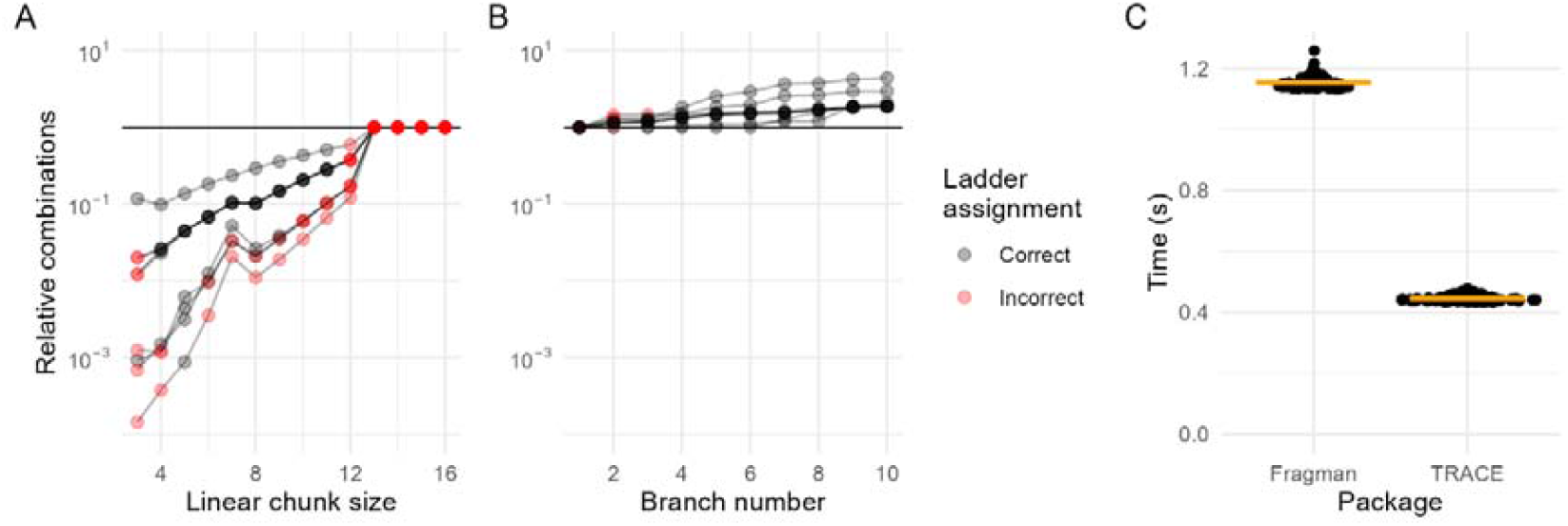
TRACE ladder assignment algorithm reduces the computational complexity of ladder-to-peak fitting. (A) Impact of linear chunk size window on the number of combinations tested relative to exhaustive search (branch number = 5). Chunk sizes >13 all have the same number of combinations because the algorithm doesn’t allow subsequent iterations to have a window of less than 4. Relative combinations are log10 transformed; lines represent the nine individual samples; black dots indicate correct final ladder assignments, red dots incorrect assignments. (B) Branch retention during iterative fitting relative to no branching (chunk size window = 5). At each chunk step, the top N branches are retained to avoid local optima. (C) Benchmarking the speed of the TRACE ladder assignment algorithm compared to Fragman R package for the same nine samples. Default settings used for each package (TRACE: chunk size window = 5, branch number = 5) with the fits plotted in supplementary 3.

Benchmarking against the R package Fragman validates this approach. In contrast to TRACE, Fragman fits polynomial models to initial peak assignments and predicts where subsequent peaks should appear. When utilized on our dataset of challenging ladders, Fragman’s assignments were not correct (Supplementary Figure 3), while also taking 2.5-fold longer (Figure 2C).

### Fragment analysis batch effects

Another challenge in fragment analysis is that repeat-containing amplicons do not migrate linearly with the ladder fragments. The spacing, in base pairs, between consecutive amplicon peaks is often smaller than the full repeat. In our mouse dataset, robust peaks within the 110–130 CAG repeat range had a median spacing of 2.85 base pairs, leading to systematic underestimation of repeat sizes when inferred directly from base-pair sizes. Across the 34 capillary electrophoresis runs, this spacing varied significantly (ANOVA, p < 0.001; Supplementary Figure 2A) with medians ranging from 2.83 to 2.87. These differences introduce batch effects that make FSA files from different fragment analysis runs not directly comparable. The gold standard for accurate repeat sizing, GeneMapper, addresses these batch effects by relying on validated size standard samples for manual calibration.

To address batch effects and/or the underestimation of repeat lengths inferred from base-pair sizes, we provide two approaches: (i) a correction factor applied across runs to bring the traces into alignment or (ii) accurate repeat sizing using validated size standard samples with known repeats. The first approach involves smoothing the signal trace and aligning the positions of the smoothed peak maxima across runs (Figure 3A), enabling the application of a batch correction factor to standardize differences (Figures 3B and 3C). This method is straightforward to implement since it requires only shared samples across runs and is useful when the accurate repeat length is not required. For example, in a cell line experiment where the average repeat change is tracked over time (McLean et al., 2024). However, it does not resolve the underestimation of repeat sizes, and comparisons between experiments without shared samples will not be possible.

**Figure 3.**
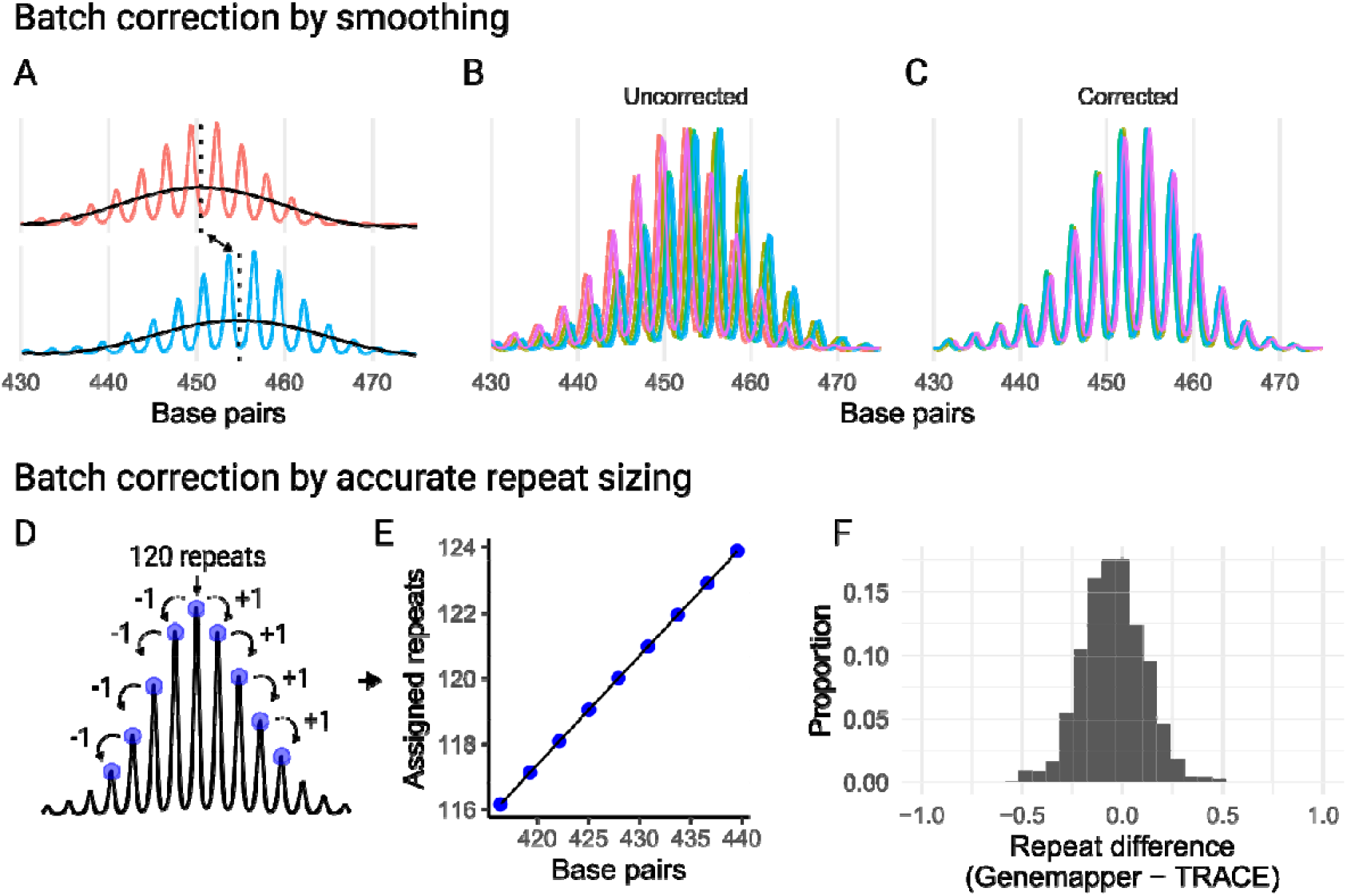
Correcting batch effects to standardize fragment analysis runs. (A) In the first approach, batch correction factors (red and blue indicate the same sample but different run) are determined by smoothing (black line) the trace and identifying maxima (vertical dotted lines), aligning traces even when they differ in their modal peaks. In this example, the top panel’s modal peak is one repeat unit smaller than that of the bottom panel when converted from base pairs. (B) The same sample from different runs is overlaid, with colors indicating different fragment analysis runs. (C) The same samples as in B, shown after applying the batch correction factor. (D) In the second approach, samples of known repeat lengths are used to assign the modal repeat length, and neighboring peaks are identified sequentially by jumping from peak to peak. (E) A linear model is created using the assigned repeats and their corresponding base-pair sizes, which is then used to predict the modal repeat length of all samples within a run. (F) The difference between the modal repeat lengths called by GeneMapper and TRACE for 872 samples across 32 runs.

### Batch correction by smoothing

The second approach accurately predicts repeat size within each run, which also corrects the batch effects. A linear model is constructed between base-pair sizes and repeats using size standards with known repeat lengths for each run. This model can then predict repeat sizes for all other samples in the run (Figures 3D and 3E). Benchmarking this method against repeat sizes called in GeneMapper, using the same size standards with known repeat length, showed high concordance, with the modal peak of 868/872 (99.5%) samples falling within the range of –0.5 to 0.5 repeat unit difference (Figure 3F). For the *HTT* CAG repeats, the linear model (R^2^ of 0.9999, base pair coefficient p < 2e-16) remained accurate up to 50 repeat units (Supplementary Figure 2C), with residual analysis highlighting increasing non-linearity beyond this range (Supplementary Figure 2D). Within the HD mouse model dataset, the percent difference in modal repeat length was negatively correlated with the base-pair spacing between peaks across runs (R^2^ = 0.44, p < 0.001; Supplementary Figure 2B). This supports the interpretation that run-to-run variation in amplicon migration rates is a major source of batch effects.

TRACE-shiny provides interactive tools that enhance the analysis and interpretation of batch correction methods. For both approaches, users can visualize the batch correction samples both before and after correction to validate the adjustments made. In the second approach, users can refine their analysis by selecting the validated repeat size of the modal peak. This is a critical feature when the modal peak shifts by chance from batch to batch. Interactivity enables users to tailor the analysis to their specific datasets, ensuring more precise and reliable repeat size predictions.

### Index assignment and metrics

In many experiments, the inherited or starting repeat length (referred to here as the index peak) is a critical reference point for downstream metrics. By default, the index peak corresponds to the modal peak of the chosen index sample. In a second sample of interest, the equivalent index peak is identified as the peak closest in repeat size to the index sample’s modal peak. In the simplest scenario, the modal peak remains the same in both samples, so the index peak is also the modal in each. In other cases, however, the modal peak of the sample of interest may have shifted due to repeat expansion or contraction; in these situations, the software still assigns the index peak based on its correspondence to the modal peak of the index sample. This allows, for example, an expanded repeat knock-in mouse liver sample to define the inherited repeat length used to assess expansion in the isolated hepatocytes (Mouro Pinto et al., 2025), or a time-zero sample to define the starting repeat length for a cell line monitored over time (Figure 4A).

**Figure 4.**
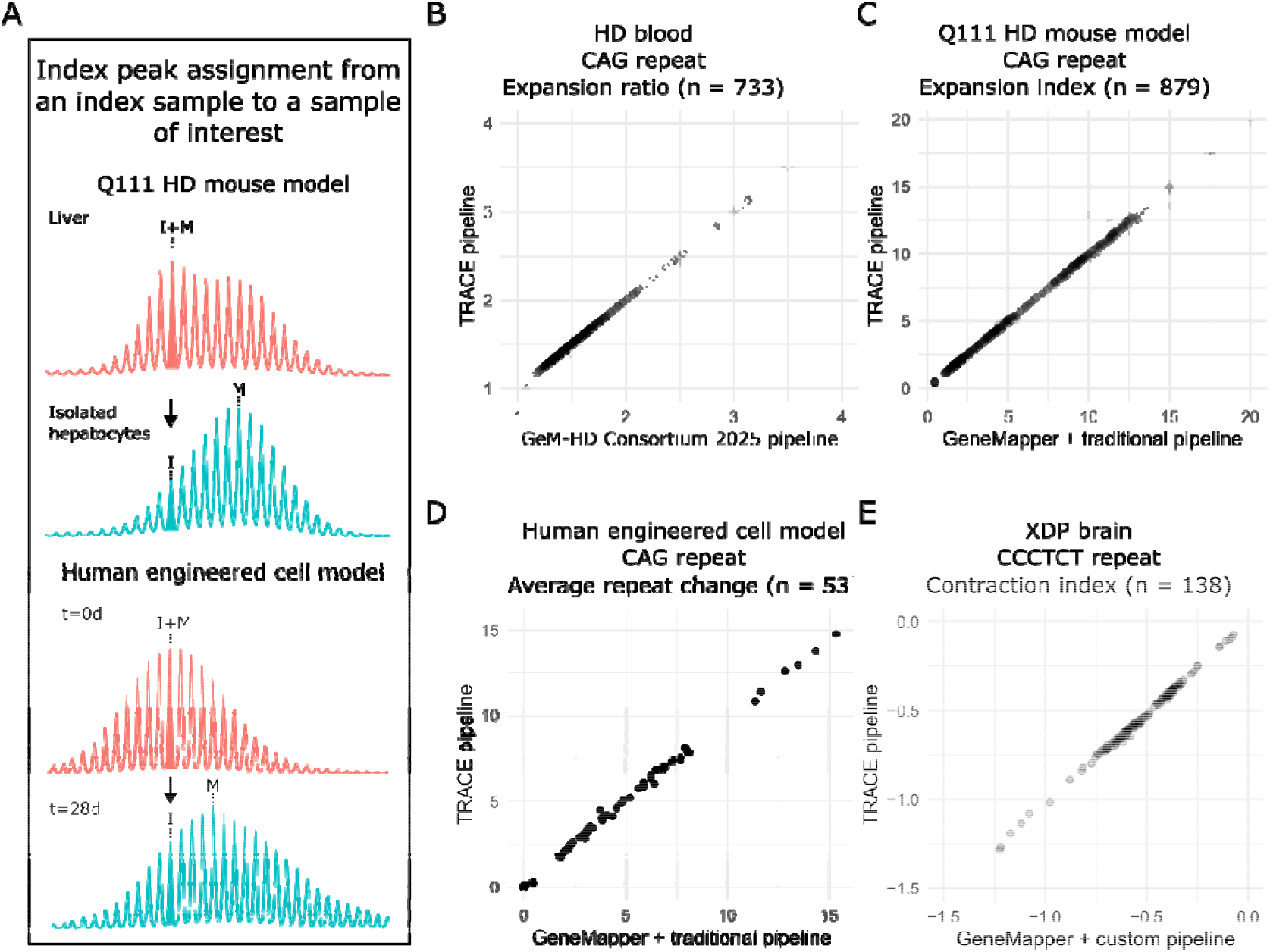
Index peak assignment and benchmarking of instability metrics. (A) Example of index peak assignment in mouse tissues (top) and cell lines (bottom). In the mouse liver sample (red), the index (I) peak corresponds to the modal (M) peak, reflecting the inherited repeat length. This index peak is then applied to the isolated hepatocyte sample (blue), where somatic expansion shifts the modal peak toward larger repeat sizes. In the cell line experiment (bottom), the modal peak of the time-zero sample (red) is used as the index peak for the 28-day sample (blue). (B) Expansion ratio in HD blood samples calculated with TRACE compared to Gem-HD Consortium data (2025). (C) Correlation of expansion index in HD mouse model samples calculated using GeneMapper with the traditional pipeline versus TRACE. (D) Average repeat change in the RPE1-AAVS1-CAG115 cell model measured with GeneMapper and the traditional pipeline versus TRACE. (E) Contraction index of CCCTCT XDP associated repeat in brain samples processed via GeneMapper and a custom pipeline compared to the values calculated by TRACE.

Finally, we benchmarked SI metrics generated by TRACE against previous pipelines. For HD blood samples, the expansion ratio calculated by TRACE was nearly identical to values from the previous custom pipeline (R^2^ = 0.999; Figure 4B). Likewise, results previously generated using GeneMapper for expansion index in HD mouse model samples (R^2^ = 0.997; Figure 4C) and average repeat gain in an *in vitro* cell model (R^2^ = 0.997; Figure 4D) showed very strong concordance with TRACE. Rare deviations in the expansion index arose because TRACE was more inclusive of peaks above threshold at the extreme of the expansion distribution, whereas the traditional pipeline excluded them once any had dropped below the threshold.

TRACE’s pipeline is versatile and accommodates diverse repeat types by incorporating amplicon size and repeat unit length into its calculations. As an example, we applied TRACE to the hexameric CCCTCT repeat associated with X-linked Dystonia-Parkinsonism (XDP) (Mejia Maza et al., 2026), which showed near identical results for the contraction index across brain tissue (R^2^ = 0.997; Figure 4E).

## DISCUSSION

Traditional methods for analysis of tandem repeat sizes in DNA by capillary electrophoresis have relied on proprietary software that required extensive manual curation and custom pipelines that vary between research groups. Here, we present an automated, open-source tool, TRACE, and associated web server TRACE-shiny. Benchmarking against data processed using the current standard analysis tool, GeneMapper, demonstrated near-identical instability metrics. We also introduce unique features such as batch correction, accurate repeat sizing, and index assignment for instability metrics calculation. Crucially, TRACE enables automated high-throughput pipelines, which are key to accelerating basic research and therapeutic development for SI. The TRACE-shiny web app requires no coding, lowering the barrier of entry to this tool.

A significant complication of fragment analysis is that repeat length cannot be directly inferred from base-pair size. We demonstrated that base pairs between consecutive peaks varied across runs and correlated with the underestimation of repeat length. This variability likely arises from differences in migration rates between repeat-containing amplicons and the ladder size standard. The resulting batch effects may influence SI metrics under specific conditions, such as (i) assigning the index peak across samples analyzed in separate runs or (ii) directly comparing metrics dependent on repeat size, such as the average repeat gain (the difference in average repeat sizes) or the difference in modal repeat size. Importantly, batch effects do not impact internally controlled metrics, such as the expansion index when the modal peak is consistently used as the index peak.

To address the underestimation of repeat length, we provide two correction approaches. The simple batch correction method is straightforward to implement and only requires shared samples across runs. This simplicity comes with the limitations that absolute repeat sizes may not be directly comparable across analyses and repeat lengths will still be underestimated. For more accurate repeat sizing, we introduced a second approach that resolves these issues but requires samples with validated repeat lengths. The effectiveness of this method depends on the quality of the reference samples, which is also true for the manual genotyping approach using GeneMapper. Ideally, the modal peaks should have clear separation in height from neighboring peaks to ensure their consistent identification. This is particularly critical for repeat lengths exceeding 100, where the signal flattens significantly due to inherent mosaicism resulting in a lack of a single predominant repeat size and/or PCR-induced errors. Overall, when carrying out SI experiments, careful planning should be taken to determine if batch correction is required and how to achieve that using these tools.

The primary goal of this tool is to generate SI metrics, which are generally relative values, rather than to accurately genotype the repeat length of samples. TRACE does not necessarily provide an accurate genotype (i.e. repeat length sizes for both alleles in human samples), so is not suitable for applications such as clinical testing. We recognize the uncertainty in the underlying capillary electrophoresis signal and therefore intentionally, as the default, do not output integer repeat units. While our repeat-calling algorithm demonstrated highly comparable allele repeat lengths to those obtained through manual GeneMapper analysis, this was limited to a single expanded allele among relatively homogenous samples. Our repeat-calling algorithm is optimized for these narrow use cases with a clearly defined expanded allele of similar size. By contrast, genotyping requires handling more complex scenarios, including variable repeat lengths, and the presence of multiple alleles that may be similar in repeat size. These factors and the importance of sequence composition of expanded repeats make sequencing approaches a more appropriate technology for genotyping.

It is possible that sequencing technologies will eventually supersede fragment analysis for quantifying SI (Ciosi et al., 2021; Maestri et al., 2024). Short-read sequencing has proven highly effective for certain applications, such as detecting instability at adult-onset repeat lengths in human blood samples (Genetic Modifiers of Huntington’s Disease, 2025). However, model systems often contain repeat tracts exceeding 100 units to enable quantification of metrics within realistic timeframes, which require long-read technologies. These long-read approaches are significantly more expensive than fragment analysis and typically require pooling large numbers of samples to bring costs into a comparable range (Ciosi et al., 2021; Maestri et al., 2024). Such designs require large-batch analyses and careful balancing of read depth to maintain accurate representations of repeat distributions. By contrast, fragment analysis remains a cost-effective and reliable method for capturing repeat distributions and continues to serve as the standard approach for calculating SI metrics across diverse systems.

Overall, TRACE and TRACE-shiny democratizes access to essential tools for SI analysis, fostering reproducibility and broadening accessibility for researchers. It is a timely resource, given the rapid growth of the repeat expansion disorder field, the increasing recognition of SI’s clinical importance, and the ongoing development of therapies targeting this process.

## Acknowledgments section

We acknowledge our community partners and all participants who provided their perspectives which informed our results.

## AUTHOR CONTRIBUTIONS

AJ: Conceptualization, Methodology, Software, Writing – review & editing. KC: Conceptualization, Methodology, Software. TG: Resources. ELO: Data curation, Resources. BPJ: Data curation. BM: Conceptualization, Software. AMM: Resources. MEM: Methodology, Funding acquisition, Writing – review & editing. RMP: Conceptualization, Resources, Methodology, Writing – review & editing. VCW: Funding acquisition, Resources, Methodology, Supervision, Writing – review & editing. JFG: Funding acquisition, Project administration, Methodology, Supervision, Writing – review & editing. ZLM: Conceptualization, Data curation, Formal analysis, Funding acquisition, Investigation, Methodology, Project administration, Resources, Software, Validation, Visualization, Writing – original draft.

## STATEMENTS AND DECLARATIONS

### Ethical considerations

Use of HD (Genetic Modifiers of Huntington’s Disease, 2025) and XDP (Mejia Maza et al., 2026) samples for benchmarking was approved by the Mass General Brigham Institutional Review Board (MGB IRB).

## DATA AVAILABILITY

TRACE is available on CRAN (https://cran.r-project.org/web/packages/trace/index.html) and TRACE-shiny is accessible at traceshiny.mgh.harvard.edu. Alternatively, a docker version is accessible through the Docker Hub under the name ‘trace/traceshiny’ or it can be installed directly from github (https://github.com/zachariahmclean/traceShiny). Data are available upon request.

