## Supplementary Figure 1 for "TRACE: Open-Source Software for Quantifying Somatic Variation of Tandem Repeats by Capillary Electrophoresis"

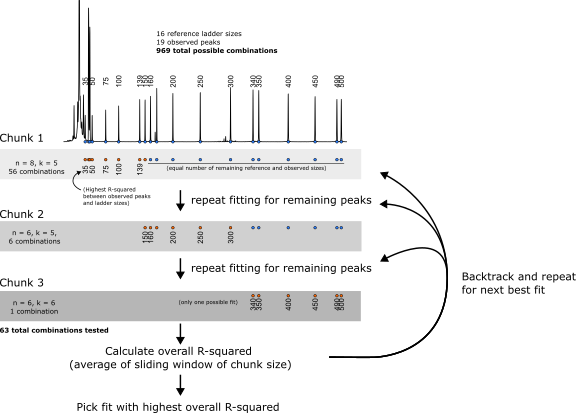

Supplementary Figure 1. Schematic of the ladder peak assignment optimization. Sixteen reference ladder sizes (text labeling the peaks in the raw ladder signal, top panel) are matched to 19 observed peaks (blue circles). Exhaustive testing of all possible assignments (969 total combinations) is computationally inefficient. To reduce complexity, assignments are fit sequentially in ordered “chunks” of the ladder. In Chunk 1 (n = 8 observed peaks, k = 5 ladder sizes), 56 possible combinations are tested, and the assignment with the highest linear R² is selected. Remaining peaks are then fit iteratively: Chunk 2 (n = 6 observed peaks, k = 5 ladder sizes, six possible combinations) and Chunk 3 (n = 6 observed peaks, k = 6 ladder sizes, one possible combination). The overall fit is calculated as the average R² across a sliding window of chunk sizes. The algorithm backtracks to test a pre-selected branching number and repeats the process. This approach reduces the number of tested combinations from 969 to 63 while maintaining accurate peak-to-ladder assignments.
