## Supplementary Figure 2 for "TRACE: Open-Source Software for Quantifying Somatic Variation of Tandem Repeats by Capillary Electrophoresis"

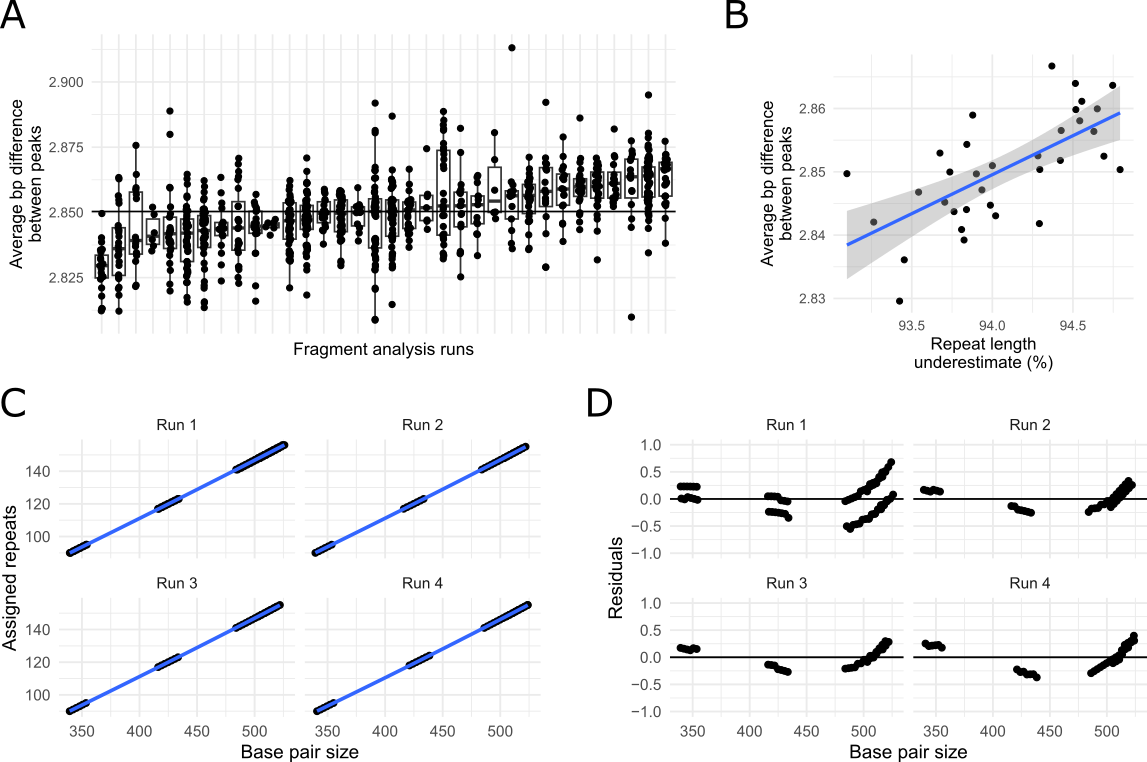

Supplementary Figure 2. Peak spacing, systematic bias, and accuracy of repeat sizing. (A) The difference between peaks in base pairs for HD knock-in Q111 mouse tissue (limited to 110–130 repeats, with a minimum of 25 peaks) is shown across individual runs, displayed along the x-axis. The blue horizontal line at 2.85 represents the median value across runs. (B) The relationship between peak spacing and repeat length underestimation. Repeat length underestimation is defined as the percent difference between the repeat length determined using the accurate repeat sizing method and that calculated directly from base pair size (i.e., the degree of repeat underestimation before correction). Each dot represents the median of these measures for a fragment analysis run. (C) The relationship between base pair size and assigned repeats using the accurate repeat sizing method across four different runs. There were three separate samples with known repeat lengths used as validated controls across the repeat length range. The close agreement across the full repeat length range reflects the high accuracy of the model. (D) Residuals from the linear model in (C), plotted on a much smaller y-axis scale (-1 to 1), show the slight non-linearity in the relationship between base pair size and assigned repeats.
