## Supplementary Figure 3 for "TRACE: Open-Source Software for Quantifying Somatic Variation of Tandem Repeats by Capillary Electrophoresis"

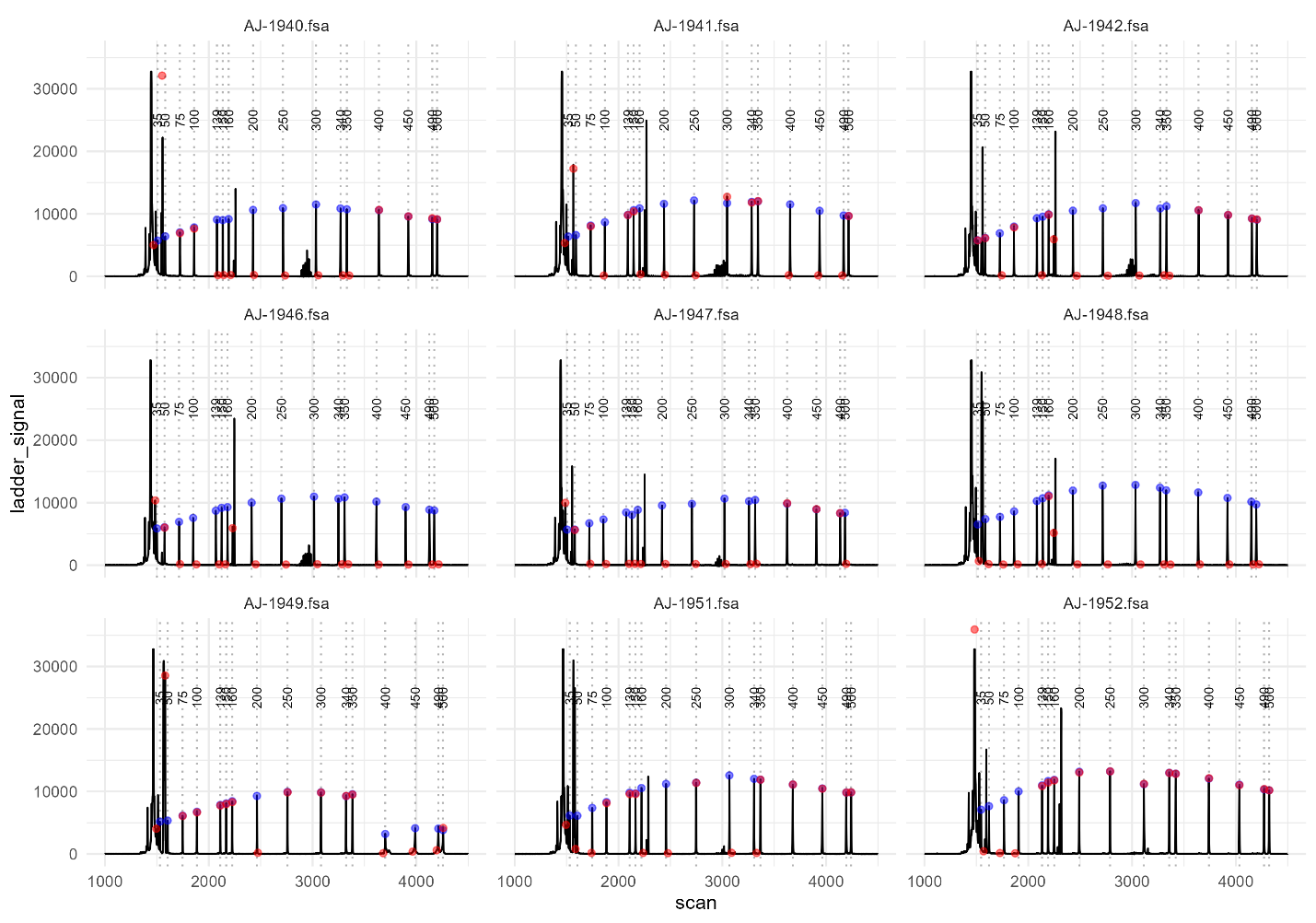

Supplementary Figure 3. Comparison the ladder assignments for trace (red) and Fragman (Blue) for the 9 test samples. The vertical dashed lines are labeled with the location and base pair size of the ladder.
